# Validation of a smartphone embedded inertial measurement unit for measuring postural stability in older adults

**DOI:** 10.1101/2020.05.29.123620

**Authors:** Friedl De Groote, Stefanie Vandevyvere, Florian Vanhevel, Jean-Jacques Orban de Xivry

## Abstract

**Background:** Identifying older adults with increased fall risk due to poor postural control on a large scale is only possible through omnipresent and low cost measuring devices such as the inertial measurement units (IMU) embedded in smartphones. However, the correlation between smartphone measures of postural stability and state-of-the-art force plate measures has never been assessed in a large sample allowing us to take into account age as a covariate.

**Research question:** How reliably can postural stability be measured with a smartphone embedded IMU in comparison to a force plate?

**Methods:** We assessed balance in 97 adults aged 50 to 90 years in four different conditions (eyes open, eyes closed, semi-tandem and dual-task) in the anterio-posterior and medio-lateral directions. We used six different parameters (root mean square and average absolute value of COP displacement, velocity and acceleration) for the force plate and two different parameters (root mean square and average absolute value of COM acceleration) for the smartphone.

**Results:** Test-retest reliability was smaller for the smartphone than for the force plate (intra class correlation) but both devices could equally well detect differences between conditions (similar Cohen’s d). Parameters from the smartphone and the force plate, with age regressed out, were moderately correlated (robust correlation coefficients of around 0.5).

**Significance:** This study comprehensively documents test-retest reliability and effect sizes for stability measures obtained with a force plate and smartphone as well as correlations between force plate and smartphone measures based on a large sample of older adults. Our large sample size allowed us to reliably determine the strength of the correlations between force plate and smartphone measures. The most important practical implication of our results is that more repetitions or longer trials are required when using a smartphone instead of a force plate to assess balance.

**Highlights:**

- We compared balance measures simultaneously obtained from a phone and a force plate
- For precision purposes, we included a large group of older participants (N=97).
- Using age as covariate, we found a moderate correlation across the two devices.
- Intra-class correlation coefficients were smaller for the smartphone.
- Balance assessment with smartphones requires longer trials compared to force plates.

## Introduction

Between 28 and 35 percent of people aged 65 and above fall at least once every year [1]. The factors that contribute to fall risk in older adults can be divided into intrinsic (e.g., age, gender, strength, vision,…) and extrinsic factors (e.g., medications, footwear, home,…) [2]. An important intrinsic factor is poor postural control, which can be defined as a complex motor skill derived from the interaction of multiple sensorimotor processes [3]. Sufficiently accurate signals from the sensory systems, effective cognitive processing and a well-functioning musculoskeletal system are key components for a good postural control [4]. It is therefore of critical importance to be able to identify people with poor postural control because they have a higher risk of falling. This can only be achieved on a large scale based on low-cost validated devices to measure postural stability outside of the laboratory. Such devices could be used by older adults and healthcare practitioners in order to assess the risk of falling.

A force plate is the state-of-the-art device to measure postural stability [5] and provides the motion of the center of pressure (COP) over time. Multiple studies have linked force plate derived measurements of the COP (e.g. increased velocity, displacement, sway area, variability and root mean square displacement) to poorer balance and higher fall-risk in older adults [6,7]. However, while some portable force plates have been developed and validated (e.g. [8]), the use of force plates outside of laboratory settings remains marginal.

In contrast, the global spread of smartphone use accompanied by the development of many health applications for mobile devices could offer a solution for providing the ageing population widespread access to objective balance assessment thanks to the embedded Inertial Measurement Unit (IMU) that they contain. Several devices with embedded IMU have been investigated for their concurrent validity for measuring postural stability [9]. Several studies have compared postural stability measures between force plates and IMUs from smartphones [10,11] (systematically reviewed in [9,12]). While most of them found a significant relationship between the outcomes of the smartphone and the force plates, these were often very variable from one condition to another. Furthermore, the small sample sizes used in previous studies prevent one to assess the strength of this relationship reliably [13] as underpowered studies tend to overestimate the size of the correlations [14]. Furthermore, several of the previous studies [9,12] did not take age as a covariate, which tends to artificially increase the reported correlation coefficients.

Therefore, the primary aim of our study is to estimate the strength of the relationship between smartphone embedded IMU and force plate measures of postural stability in a large group (~100) older adults across four conditions (eyes open, eyes closed, dual-task and semi tandem condition).

## Methods

### Participants

99 older adults between 50 and 90 years old participated to this study. Two people were excluded from the analysis (one person who was incapable of completing the cognitive task and one person with an acute loss of sensibility and strength in the lower legs) leaving us with 97 participants (42 males and 55 females). Participants were recruited through local advertisements at typical gathering places for older adults, through mailings to organisations for older adults, and within the social network of the research team. The exclusion criteria were: 1) being younger than 50 or older than 90 years; 2) being diagnosed with severe neurological, medical, or psychiatric diseases; 3) being incapable of understanding the test instructions; 4) depending on a wheelchair or other walking aids, except for a walking stick. The study was approved by the local ethics committee of UZ/KU Leuven (S61583). All participants signed an informed consent form in accordance with the Declaration of Helsinki.

### Materials

The BTrackS Balance Plate (BBP, BTrackS™ Balance Plate, California, USA) was used for measuring COP movement [8]. This force plate contains force transducers beneath each corner of the rectangular plate (40 cm x 60 cm). The BBP measurements were registered with a sample frequency of 25 Hz. The BBP was placed on a stable surface. The BtrackS Explore Balance software was used for analysing the COP-measurement data.

Smartphone measurements were made using a Samsung Galaxy S7 smartphone (Samsung, Seoul, Korea) attached to the subject’s waist near the body’s center of mass (COM) location. The IMU of the Samsung Galaxy S7 smartphone consists of an accelerometer and gyroscope. IMU-data were collected by a custom-made smartphone application developed in the Faculty of Movement and Rehabilitation Sciences at KU Leuven. It registered the linear and rotational acceleration of the smartphone with a mean sample frequency of 500 Hz (range 499 Hz-509 Hz).

### Procedure

We used a cross-sectional study design to examine the concurrent validity of the Samsung Galaxy S7 smartphone in static balance conditions. The measurements were performed at the Faculty of Movement and Rehabilitation Sciences or at the participant’s home. First, the participant filled out a fall risk assessment. Next, the participant was instructed to stand barefoot on the force plate (1 meter in front of the wall) in a comfortable posture, both hands positioned at hip level. The position of both feet was tape-marked to maintain the same position on the force plate for the different conditions. The belt with the smartphone was positioned on the participant’s back by the instructor at the level of the second sacral vertebra, near the body COM (Fig. 1). During the entire test procedure, the instructor stood behind the participant to guarantee his/her safety. Balance was assessed in four conditions: 1) bipodal stance with eyes open (BSEO), 2) bipodal stance with eyes closed (BSEC), 3) a dual-task consisting of bipodal stance with eyes open and a cognitive task (CTEO), and 4) semi-tandem stance with eyes open (STEO). During the dual task, participants were attributed a random number between 100 and 120, and were asked to count down aloud in steps of 7. Each balance test lasted 35 seconds and was repeated four times. Every first trial from each balance test was categorised as a familiarisation trial and was not included in our analysis. Participants were instructed to stand as still as possible while looking at a standardised fixation point on the wall, 1 meter in front of them. At the end of every test condition, the participant got off the force plate and had the opportunity to rest while the instructor changed the set-up for the next test. For the fourth balance test, the instructor turned the force plate 90° to allow the participant to place both feet on the plate in semi-tandem stance. If the patient lost his/her balance during the trial, the trial was repeated.

**Figure 1:**
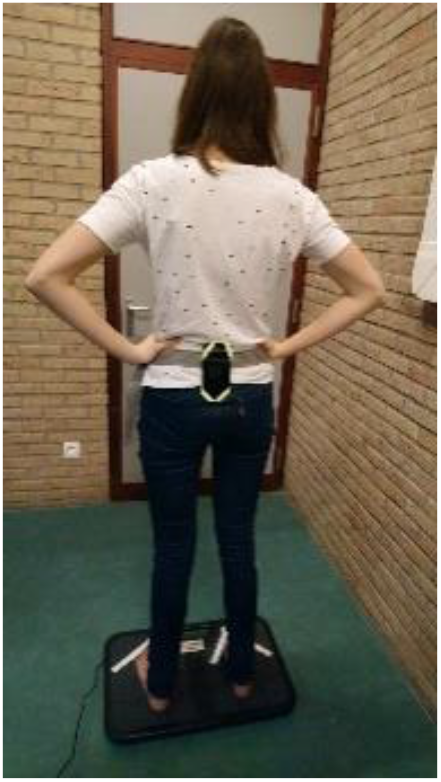
Test setup. Participants stood barefoot on the force plate (1 meter in front of the wall) in a comfortable posture, both hands positioned at hip level. The smartphone was attached to the participant’s back at the level of the second sacral vertebra, near the body center of mass.

### Data analysis

Only data collected during the last 30 seconds of each 35 seconds trial was analyzed. COP position measurements from the force plate were processed with a Savitzky-Goley filter (5th polynomial order and frame length of 7) to obtain filtered position, velocity and acceleration signals. The average value was subtracted from each position, velocity and acceleration signal. This gave us three signals in two different directions (anterior-posterior and medio-lateral). For each of these signals, we computed the root-mean-square (RMS) and the mean absolute value (Mean) resulting in 12 force plate outcome measures (2 directions x 3 derivatives x 2 analysis types). We only used smartphone acceleration data in the anterior-posterior and medio-lateral directions (smartphone-based reference). We first subtracted the mean from the signal and then computed a weighted RMS and mean absolute value as the sampling frequency was variable, resulting in four smartphone outcome measures. Data processing was done in Matlab (R2018b, Mathworks, Natick, Massachusetts, USA). Raw data and analysis scripts can be accessed at https://osf.io/vpd79/?view_only=07b131728d934f5e9948fb545a7810e2

The selection of the above mentioned parameters was based on published parameters from a study with a similar experimental set-up for measuring the concurrent validity of a mobile device according to a force plate to measure postural stability and fall risk in elderly [10].

### Statistical analysis

All statistical analyses were performed in R [15] based on the average outcome measures over the three trials, except when analyzing intra-class correlations. We used a robust correlation method (80% winsorized correlation) using the wincor function from the WRS2 package [16].

In the first analysis, we correlated all parameters with age. The confidence interval of these coefficients was computed via a bootstrap procedure. In the second analysis, we considered the eye opens condition as reference and computed the magnitude of the difference for the three other conditions by means of Cohen’s d (within-subject difference between the two conditions divided by its standard deviation). Confidence intervals were obtained by the ci.sm function from the MBESS package [17]. In the third analysis, we computed the intra-class correlation coefficient across the three trials for each participant with the test-retest function (trt) from the Relfeas package [18].

Because all parameters were correlated with age, we regressed age out of all ensuing correlation analyses. In the fourth analysis, we computed the winsorized correlation between all parameters from the force plate with all parameters from the smartphone. We did not correct for multiple comparisons, as we were not interested in the significance of these correlations but in their magnitude. The confidence interval of these coefficients was computed via a bootstrap procedure. Finally, we performed a factor analysis on all signals (partialling out the effect of age) from the force plate and the smartphone separately. In this factor analysis, we considered that postural stability was the only latent factor. We used the fa function from the psych package [19] with oblimin rotation. The factors were obtained by maximum likelihood factoring method and the factor scores by the regression method. Factor scores for the force plate and smartphone obtained separately were then correlated with a winsorized correlation. Scripts for statistical analysis can be accessed at https://osf.io/vpd79/?view_only=07b131728d934f5e9948fb545a7810e2

## Results

### Demographics

In this study, we investigated whether a smartphone IMU was sufficiently reliable to assess balance. To answer this question, we assessed balance in 97 older adults with a force plate to measure the COP and with a smartphone fixed on the bottom of their back. Our participants were between 50 and 90 years old. Few of them required a walking aid, a portion of them did not have maximal walking confidence, and 13 of them had already experienced falls (Fig. 2, data derived from fall risk assessment).

**Figure 2:**
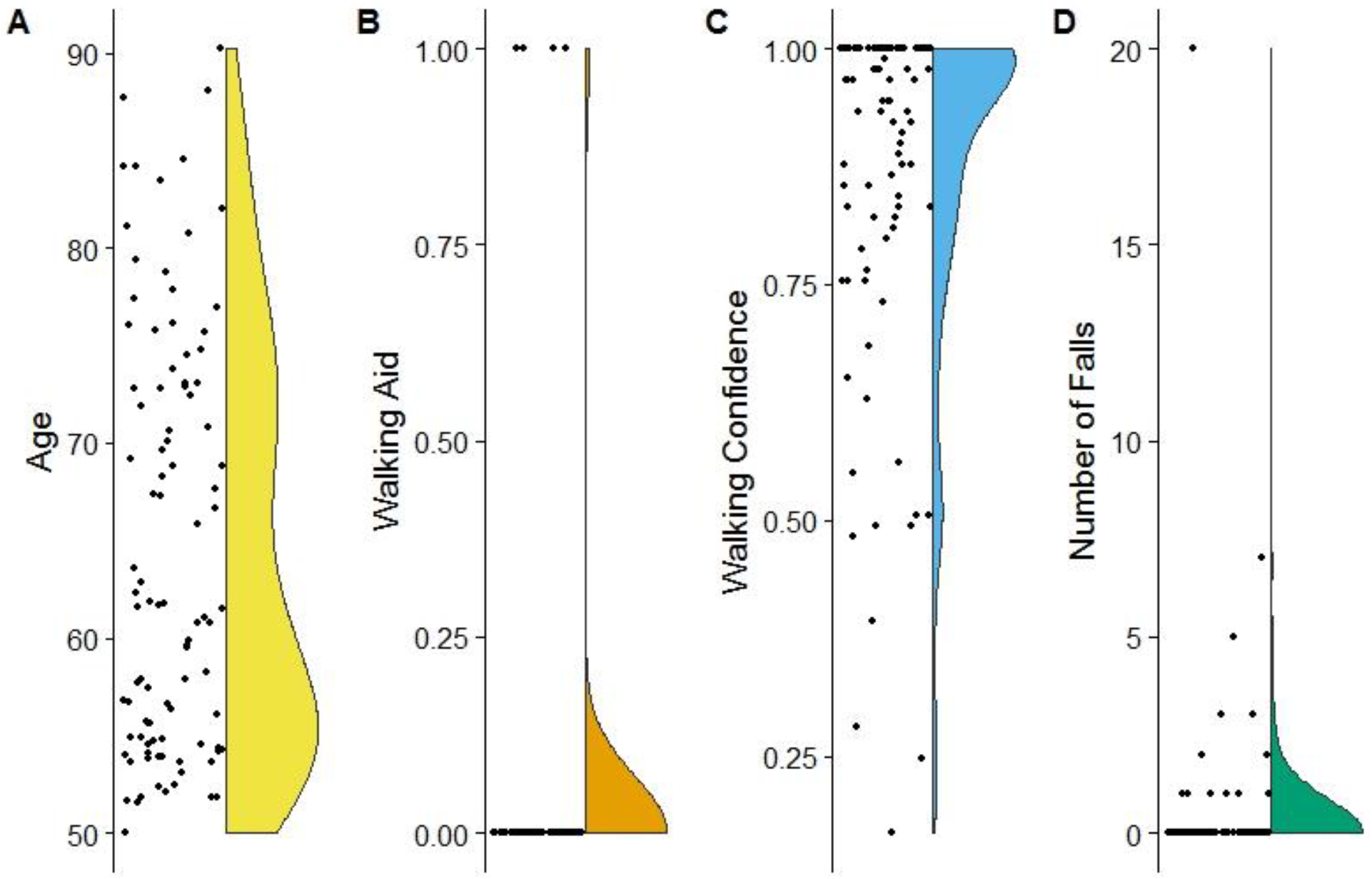
Description of participants. (individual points + smoothed distribution): A) age of the participants; B) using a walking-aid (yes=1, no=0); C) walking confidence as measured by a visual scale (high=1, low =0); D) self-reported number of falls in the last year.

### Effect of age on balance

All outcome parameters were correlated with the age of the participants with correlation coefficients around 0.5 for force plate parameters derived from COP velocities and accelerations while these correlations were around 0.35 for the smartphone parameters, showing that the force-plate might be slightly more sensitive to measure age-related changes in postural-sway than the smartphone (Fig. 3A). In other words, while one needs 30 participants to detect an effect of age with the force plate, one needs 61 participants with the smartphone (80% power, significance level of 0.05).

**Figure 3.**
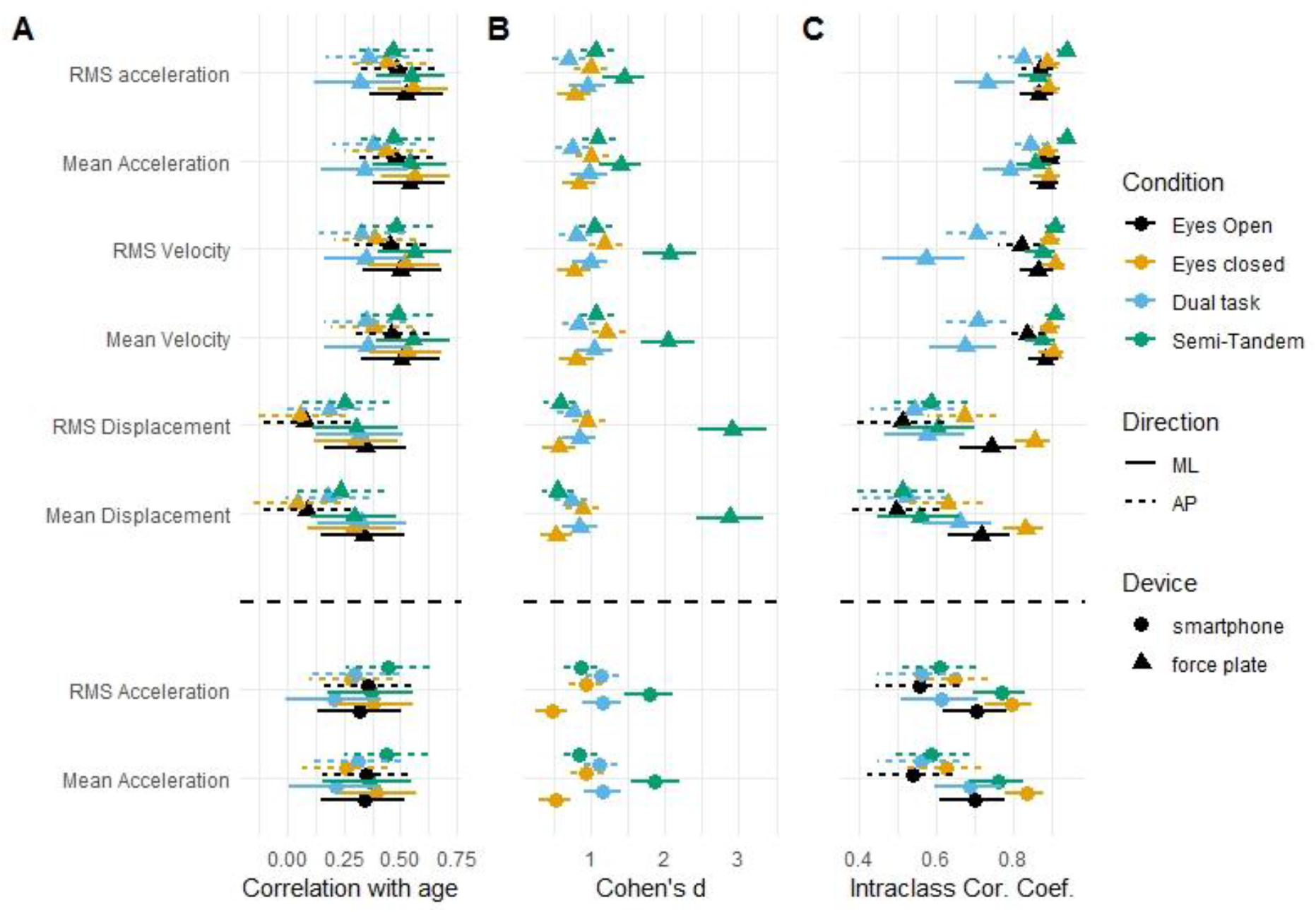
A) Sensitivity of the different measurement to age-related changes in balance (symbol and error bar represent mean and confidence interval of the winsorized correlation coefficient) for the different directions (medio-lateral: ML and anterio-posterior: AP), conditions, and devices. B) Sensitivity of the different measurements to the difference in postural sway in the different experimental conditions relative to the eyes open condition (symbol and error bar represents mean Cohen’s d and confidence interval). C) Reliability of the different measurements over the three test trials as assessed by the intra-class correlation coefficient (ICC) (symbol and error bar represent mean ICC and confidence interval). All variables were first corrected for age.

### Sensitivity of outcome parameters to different conditions

Force plate and smartphone derived outcome parameters captured differences between the different conditions and the eyes open condition (which was taken as reference) equally well with effect sizes around 1 (Fig. 3B). Effect sizes were largest for the semi-tandem condition in the medio-lateral direction but varied quite a lot among the different force plate outcome parameters.

### Reliability of outcome parameters

The test-retest reliability of the outcome parameters across the three repetitions differed between conditions (Fig. 3C). In the dual-task condition, measures derived from COP position and velocity were more variable than in the other conditions, reflecting that the cognitive load imposed by dual-tasking can be highly variable from trial to trial. When not considering the dual-task condition, the test-retest reliability of the smartphone outcome parameters was moderate (0.5-0.75) to good (0.75-0.9) while the test-retest reliability of the force plate outcome parameters was excellent (>0.9).

### Correlation between smartphone and force plate outcomes

It is of high importance to know whether these two devices measure postural sway in the same way and whether someone who would be characterized as having bad postural stability would be so with both devices. Therefore, we correlated the outcome parameters of the smartphone with those of the force plates in all conditions and directions separately.

It was important to use robust correlation methods as a few points were away from the main populations and could drive spuriously high correlations as illustrated in Fig. 4 for the eyes open and semi-tandem conditions in the medio-lateral direction. Walking confidence was not highly linked to high postural sway as darker points are scattered along both axes and not on the top right of the graphs.

**Figure 4:**
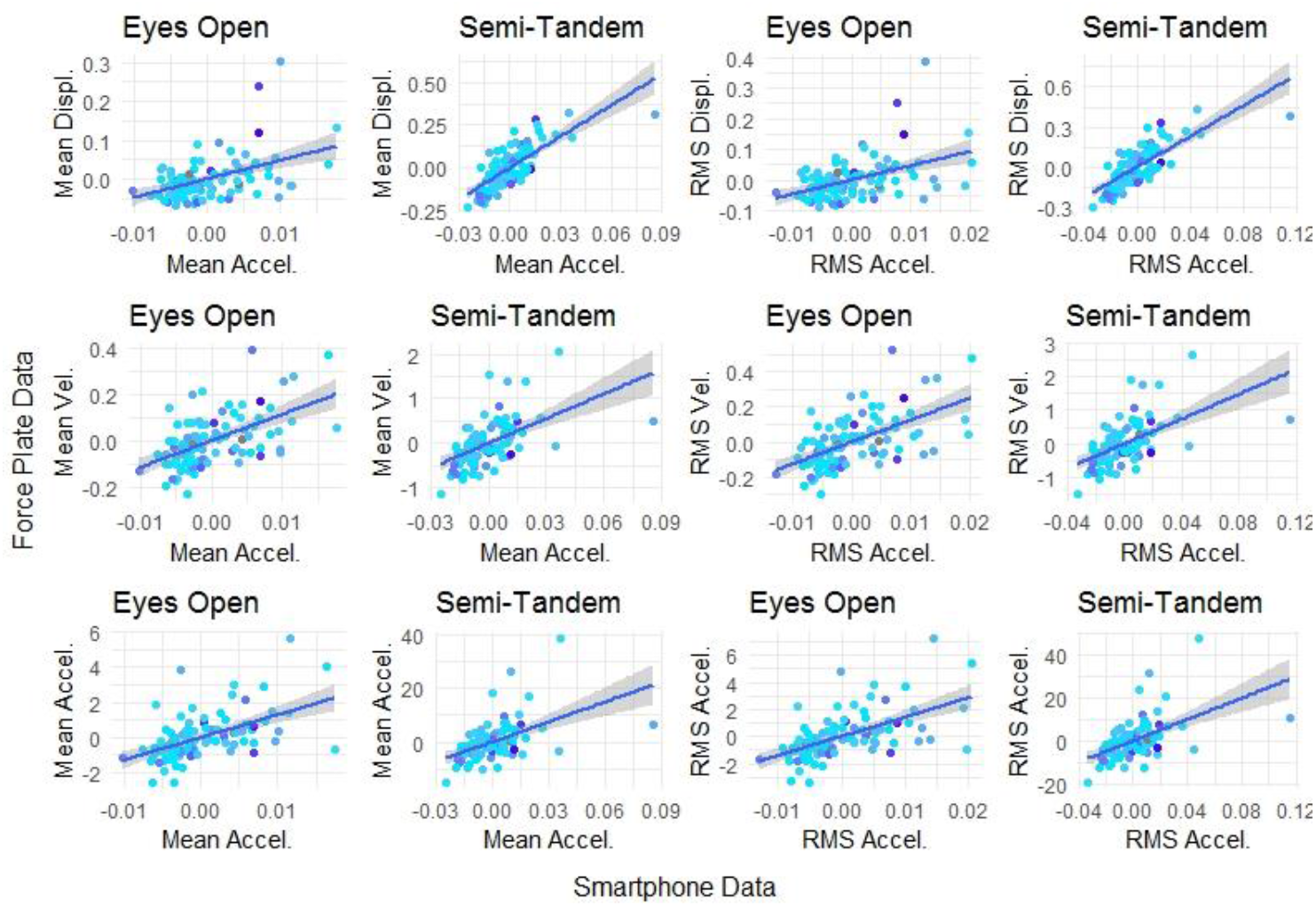
Correlation between outcome parameters for the smartphone (horizontal axis) and for the force plate (vertical axis) for the eyes open (EO) and semi-tandem (ST) conditions. Outcome parameters for the force plate vary along the rows (displacement - Disp, velocity - Vel and acceleration - Acc). First two columns focus on the mean absolute value, last two on the root-mean-square value. The color code represents walking confidence with high walking confidence in light blue and low walking confidence in dark blue. Blue line represents the regression line for the normal regression with the grey area representing its 95% confidence interval

Outcome parameters from the force plate and the smartphone are correlated but the extent of these winsorized correlation accounting for age depends on conditions and directions with a range from 0.14 to 0.82 (Fig. 5). Furthermore, the correlations are often larger in more challenging conditions (medio-lateral direction in semi-tandem, anterior-posterior direction during eyes closed) than in the easier conditions.

**Figure 5:**
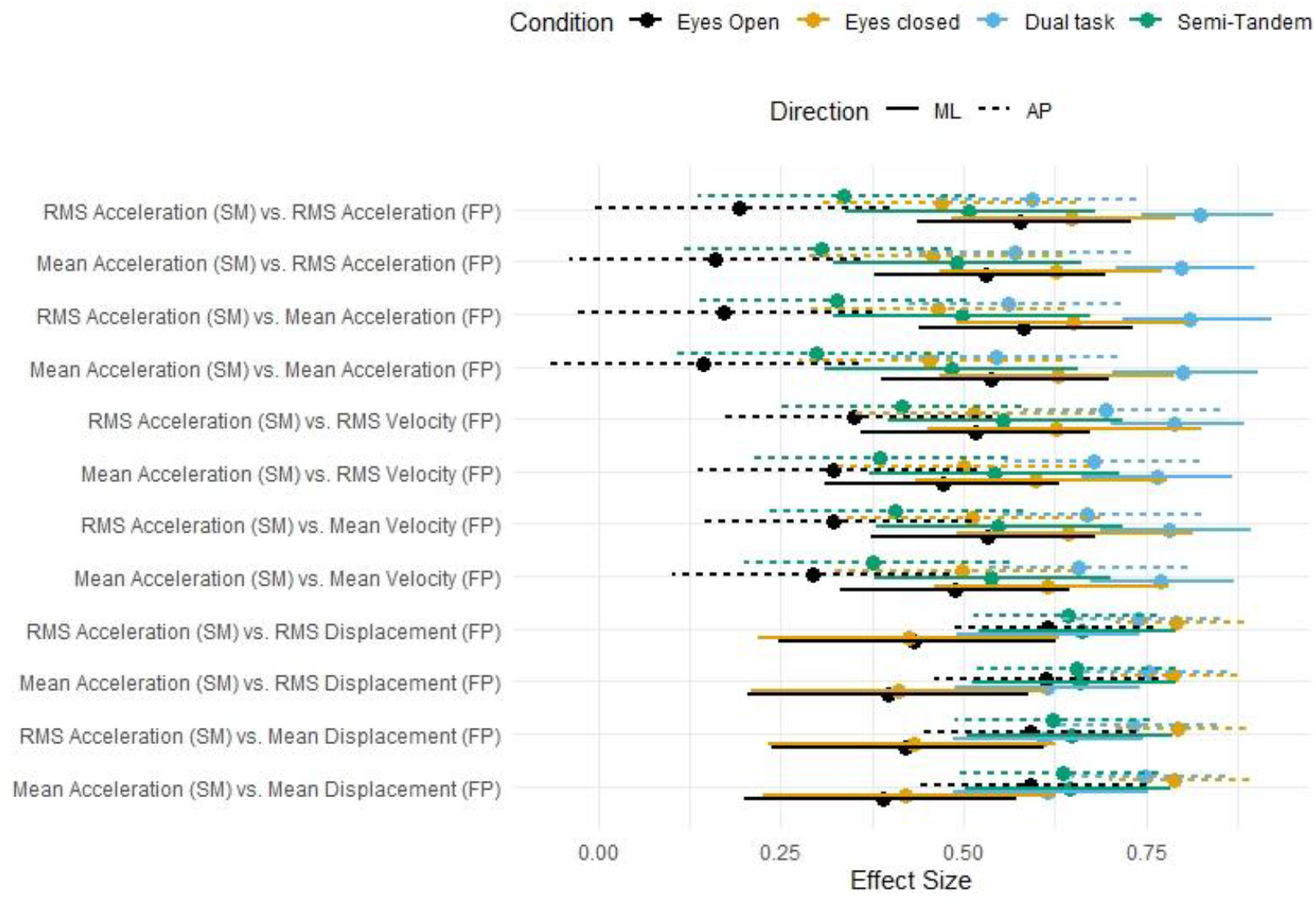
Correlation between force plate and smartphone parameters. Mean and bootstrapped confidence intervals for all correlations between every pair of outcome parameters from the force plate (FP) and smartphone (SM) in all directions and all conditions. Correlation coefficients were obtained via winsorized correlations accounting for the effect of age.

Yet, this correlation range does not allow us to assess how the measures of postural sway with the smartphone and with the force plate are related. Therefore, we decided to use factor analysis (with a single latent factor) on the force-plate and smartphone data (partialling out the effect of age) separately in order to extract the latent factor linked to the control of balance with both devices. The factor scores with a single latent variable, postural stability, obtained from either all force plate or all smartphone measures were correlated with a winsorized correlation coefficient of 0.58 (CI=[0.42, 0.75], Fig. 6).

**Figure 6:**
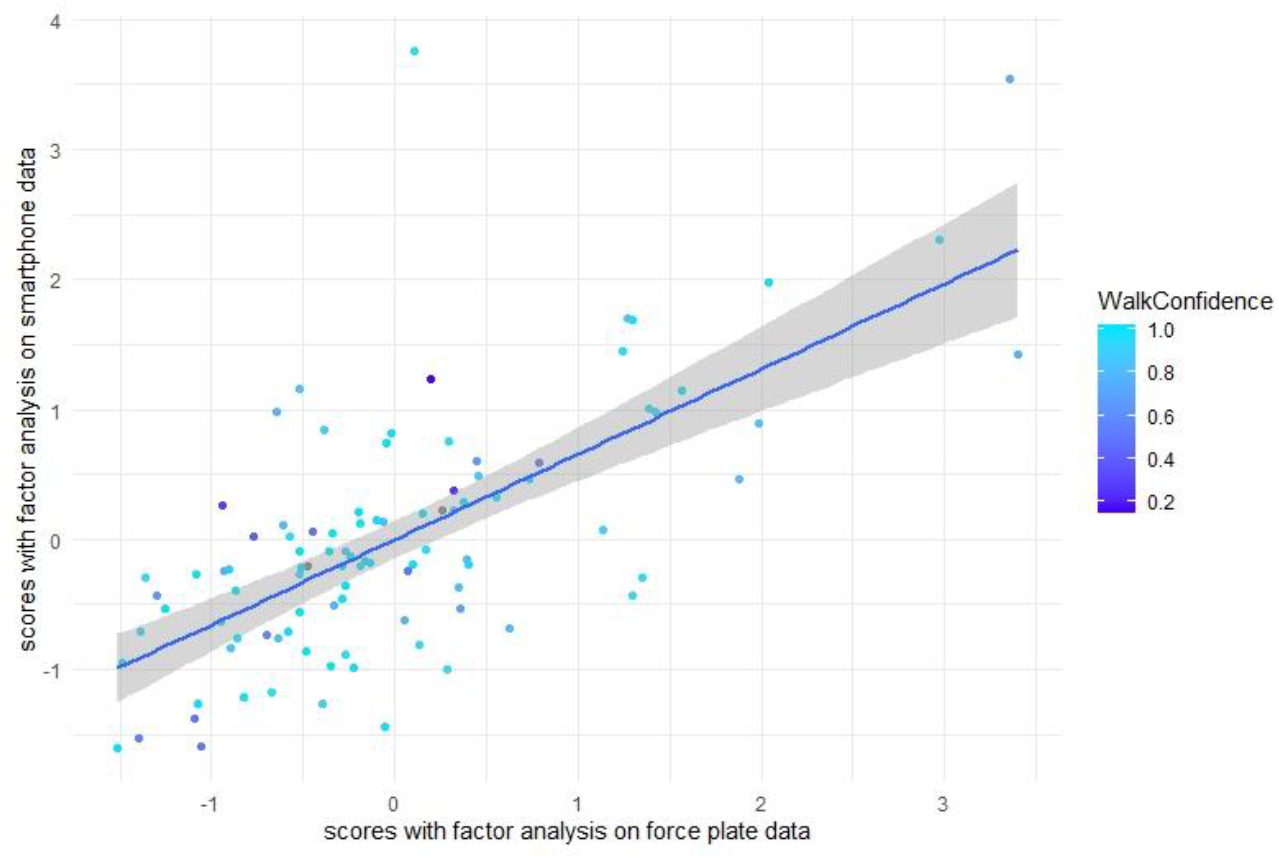
Correlation between factor score obtained from force plate outcomes and factor score obtained from the smartphone outcomes accounting for the effect of age. Color of the points provides info on the walking confidence of the different participants. Blue line represents the regression line for the normal regression with the grey area representing its 95% confidence interval.

## Discussion

The primary aim of this study was to investigate the extent to which the IMU embedded in a smartphone could be a valid instrument for measuring postural stability in older adults. While previous studies suggested that there was a link between measures of postural stability obtained by a force plate and IMU, none of them was able to assess the magnitude of this relationship with sufficient accuracy because of small samples and the presence of covariates such as age.

Our study brings a few important messages about the measurement of postural instability with an IMU compared to with a force plate. First, the correlation of postural stability parameters with age can be detected with both IMUs and force plates but the force plate measures derived from COP position and velocity appear to be slightly more sensitive. Second, differences across conditions could be detected equally well with both systems. This is in line with previous studies showing that the accelerometer could distinguish between balance tasks [20,21]. Third, the test-retest reliability was larger for the force plate than for the smartphone but was also dependent on the direction of the sway and on the condition with the dual-task condition being the least reliable. Finally, the robust correlation coefficient between the parameters from the smartphone and the force plate (with age regressed out) was very variable across conditions and parameters but was mostly of moderate magnitude (around 0.4-0.5). The factor analysis, which takes all conditions, directions and parameters into account, was moderately correlated as well (r=0.58, CI=[0.42, 0.75]).

While these results support previous conclusions that smartphone embedded IMUs could be used to assess postural instability [10], our results also highlight that they are not as good as force plates. IMUs have several advantages including their small size and price. However, they are not as precise as the force plate and they are also more variable from trial-to-trial, given their lower intra-class correlation coefficient. These observations cast doubts on the fact that smartphones could fully replace force plates in the future.

Future use of IMUs to measure postural stability should use longer trials because this could potentially increase the reliability of the measures. Yet, one also needs to pay attention to fatigue effects when trials are becoming longer.

The location of the smartphone on the body should be investigated further. In Hsieh et al [10], participants held the smartphone vertically against their chest while we attached it to the subject’s lower back, which should be close to the body COM, therefore yielding easier to interpret measures.

The reported correlations could be underestimated as the smartphone and the force plate measure different signals. While the force plate measures COP location, the smartphone provides accelerations of a point close to the body’s COM. COP and COM positions are related through the body dynamics but this relation is not linear [3]. Moreover, literature shows that different subjects may use different correcting strategies with different movements in the ankle, knee and hip joints depending on e.g. the age and fear of falling, which is expected to negatively affect the correlation between both measures [22].

The semi-tandem condition is the optimal choice for assessing balance with a smartphone as this combines a relatively high ICC (Fig. 3) and a moderate correlation with the force-plate measures (Fig. 5) across both directions. With the smartphone, test-retest reliability was higher in the eyes closed and semi-tandem conditions in the medio-lateral direction and the correlation with the force plate outcome was consistently high (high values and smallest confidence interval, Fig. 5) in the ML and AP direction in the semi-tandem condition. Furthermore, the correlation between COP displacement and COM acceleration was the most consistent in the semi-tandem condition in both directions. Therefore, the semi-tandem condition seems to be the best one to measure postural stability in a reliable manner. This contrasts with the fact that the AP condition during bipodal stance has been considered as the most sensitive condition for reflecting the effect of age on postural stability when measured with a force plate [23]. Our results show that the outcomes are equally sensitive to age across several conditions (Fig.3).

Our study bears several limitations. First, the placement of the smartphone was not always optimal due to the body curvature of the participant, and hence the measurement axis of the IMU was not always perfectly aligned with the anterio-posterior direction. Integration of gyroscope data from the smartphone embedded IMU might be used to correct this aberrant position in the data analysis. The testing was done at several locations (at people’s home or at the Faculty of Movement and Rehabilitation Sciences). Different test environments and ambient noise may have influenced the results. Third, our population sample was rather homogenous, consisting only of healthy individuals. Therefore, the applicability of our results to other populations (e.g., younger people, individuals with balance disorders, etc.) should be investigated. Fourth, we did not use a clinical scale, e.g. Berg’s Balance Scale, to assess our participants. Future studies should investigate whether clinical screening on the risk of falling could be complemented by quantitative assessment with IMUs in order to detect elderly with a higher fall risk.

## Conclusion

In conclusion, a smartphone embedded IMU could be used to assess balance in older adults but trials need to be longer or repeated more often as compared to force plate measurements in order to deal with the lower test-retest reliability.

## Notes

https://osf.io/vpd79/?view_only=07b131728d934f5e9948fb545a7810e2

### Competing Interest Statement

The authors have declared no competing interest.

